# Insights into the trafficking of human leukocytes to colostrum evidences a modulation of the B lymphocyte compartment in obesity

**DOI:** 10.1101/2021.11.19.469333

**Authors:** Raúl Piñeiro-Salvador, Eduardo Vazquez-Garza, José Antonio Cruz-Cardenas, Cuauhtémoc Licona-Cassani, Gerardo de Jesús García-Rivas, Jorge Moreno-Vásquez, Mario René Alcorta-García, Victor Javier Lara-Diaz, Marion E. G. Brunck

**Affiliations:** Tecnologico de Monterrey, School of Engineering and Sciences, Av. Eugenio Garza Sada 2501 Sur, Tecnologico, 64849, Monterrey, Nuevo Leon, Mexico; Tecnologico de Monterrey, Escuela de Medicina y Ciencias de la Salud, Ave. Morones Prieto 3000 Poniente, Col. Doctores, 64710, Monterrey, Nuevo Leon, Mexico

**Keywords:** Colostrum, Leukocytes, Obesity, Flow cytometry

## Abstract

Breastmilk is a dynamic fluid which initial goal is to provide the most adapted nutrition to the neonate. Additional functions have been recently attributed to breastmilk, with the evidence of a specific microbiota and the presence of a variety of components of the immune system, such as cytokines and leukocytes. The composition of breastmilk varies through time, according to the health status of mother and child, and altogether contributes to future health of the infant. Obesity is a rising condition worldwide, that creates a state of systemic, chronic inflammation including leukocytosis. Here, we asked whether colostrum, the milk produced within the first 48 h post-partum, would contain a distinct leukocyte composition depending on the body mass index (BMI) of the mother. We applied a panel of 6 antibodies plus viability marker to the peripheral blood and colostrum obtained from obese (BMI > 30) and lean (BMI < 25) mothers to characterize 10 major leukocyte subpopulations using flow cytometry. While lymphoid cells were otherwise unaffected by their tissue of origin, the phenotypes of granulocyte and monocyte populations significantly contrasted between blood and colostrum, including variations in morphology and surface expression of CD45 and CD16. These differences recapitulated across groups, which suggests a generalized cell-specific phenotype alteration caused by trafficking to colostrum. The B lymphocyte compartment was significantly reduced in obese colostrum and these cells did not exhibit enhanced CD16 shedding in this tissue, unlike B lymphocytes from lean mothers’ colostrum. This is the first exhaustive characterization of major leukocyte subsets in obese mothers’ colostrum, and the first report of leukocyte subpopulations from Latin-American women’s colostrum. This pioneering study is a steppingstone to further investigate active immunity in human breastmilk.

## 1. INTRODUCTION

Human breastmilk has been historically regarded primarily as a source of nutrition for infants. Recent studies have evidenced that breastmilk is actually a complex and dynamic tissue that provides newborns with components involved with functions beyond nutrition, such as the breastmilk microbiota and mother-derived cytokines and leukocytes^1^. As a well-studied consequence, infants, born with an immature immune system, benefit from the passive transfer of antibodies from their mother through breastfeeding. This includes immunologically relevant concentrations of immunoglobulin in breastmilk over a long period of time, and vaccine-induced antigen-specific IgA and IgG into breastmilk 2-6 weeks post-vaccination^2,3^. As a consequence, unvaccinated infants benefit from antibody-mediated protection against infectious diseases, in addition to training of their immature immune system by exposure to these components^4^.

Much less is known about the regulation and biological relevance of mother-derived leukocytes in human breastmilk, despite speculations of active immunity transfer^5^. To understand the implications of leukocytes in breastmilk, the causes of initial recruitment to this tissue in homeostasis should be elucidated. However, to the best of our knowledge, mechanisms of recruitment of leukocytes to breastmilk have not been investigated to date. The leukocyte composition of breastmilk greatly varies within a single feed, through lactation stage, with pregnancy term, and with the health status of mother and infant, and colostrum is the most leukocyte-rich type of breastmilk^6,7^. Recent work in mice suggests that a more alkaline pH of newborns stomach, combined with loose cellular junctions in the GI tract, allow mother-derived cells to translocate to various organs of suckling pups^8,9^.

While providing infants with immunity to infectious agents is beneficial, the transfer of immune factors through breastmilk may also promote the development of autoimmune conditions^10,11^. Obesity rates are increasing worldwide. Mexico is an unfortunate global leader in obesity prevalence with > 40% women presenting a Body Mass Index (BMI) > 30, and increasing rates of obesity in the younger populations^12,13^. Obesity is now regarded as a state of low-grade systemic inflammation where larger adipocytes secrete proinflammatory mediators and recruit leukocytes^14,15^. Obese individuals have altered peripheral blood cell counts with increased risks of leukocytosis, and modulations in the phenotype of lymphocyte subpopulations^16,17^. In breastmilk, obesity impacts the macro- and micro-nutrients^18^. We recently evidenced that colostrum from obese mothers suffered significant adjustments in α- and β-indices of microbiota diversity^19^. However, the consequences of obesity on various breastmilk leukocyte populations have not been reported to date^20^.

Here we measure the frequency of 10 major leukocyte subpopulations in peripheral blood and colostrum obtained from lean and obese mothers within 48 h post-partum, by applying an optimized 7-colour panel for flow cytometry^7,21^. We identified considerable and cell-specific phenotypic alterations of all leukocyte subtypes investigated in colostrum, which inform for the first time on regulated processes of leukocyte trafficking from human blood to colostrum. We also evidenced a reshaping of the colostrum B lymphocytes compartment in obesity, that is significantly reduced and exhibits a decreased ability to shed CD16 compared to lean mothers’ colostrum B cells. This pioneering work contributes to advancing our understanding of breastmilk-mediated contributions to early life development and its possible influence on chronic diseases predispositions.

## 2. RESULTS

### 2.1 The experimental design highlights significant difference between study groups

We processed blood and colostrum samples from a total of 21 volunteers in the lean mothers’ group and 22 volunteers in the obese mothers’ group. Post-acquisition quality filters restricted the final analysis to 21 blood samples and 17 colostrum samples in the lean mothers’ group, and 20 blood samples and 11 colostrum samples in the obese mothers’ group. A summary of the clinical variables of each study group is presented in Table 1. As per study design, the average BMI was significantly larger in the obese mothers’ cohort (33.6 vs. 22.5). There was no difference in gestational age at term, mode of delivery (C-section or vaginal), or gender of infant between groups. The average volume of collected colostrum was significantly smaller in the obese mothers’ cohort (1.42 ml vs 2.11 ml). Difficulties in initiating lactation have been reported for obese mothers, which are in part explained by a decreased prolactin production as a response to infants suckling early post-partum in obese mothers ^22-24^.

**Table 1.** Summary of demographics and clinical parameters of study participants

| Variable | Cohort |  | *p* |
| --- | --- | --- | --- |
|  | Lean | Obese |  |
| N total | 21 | 20 |  |
| Maternal BMI (kg/m<sup>2</sup>), mean (SD) | 22.5 (1.5) | 33.6 (3.9) | **<0.001** |
| Infant gender, N females (% total) | 9 (42.8%) | 8 (40%) | 1 |
| Gestational age, weeks (SD) | 39.3 (0.9) | 38.7 (1.1) | 0.09 |
| Delivery type, N V (%) | 15 (71.4%) | 13 (65%) | 0.73 |
| Volume of colostrum obtained (mL), mean (SD) | 2.11 (0.93) | 1.42 (0.74) | **0.012** |

BMI: body mass index, V: vaginal births

### 2.2 The majority of leukocyte proportions are significantly regulated between blood and colostrum

Colostrum samples were enriched for cellular fractions before staining and flow cytometry analysis. Mean viability of colostrum enriched cells, as assessed visually by trypan blue exclusion, was > 90% in both cohorts (median: 90.6%, range [68.6-98.9]). This is similar to what has been described previously using the same methodology for viability assessment^7^. Only recently have flow cytometry analyses of breastmilk cells incorporated a viability stain in their panel^25,26^. Here, the addition of propidium iodide as a viability stain is an optimization of a panel previously applied to human milk samples, which allowed to eliminate on average 6.06% and 6.30% of dead cells from downstream analyses from fresh colostrum single events collected from lean and obese groups, respectively (Fig. 1). The simultaneous determination of sample viability, together with surface markers measurements, excluded non-specific staining of dead cells from the final analysis. This could impact significantly final proportions as 8 out of 10 subpopulations identified had <4% frequency amongst colostrum leukocytes (Table 2).

**Fig 1.**
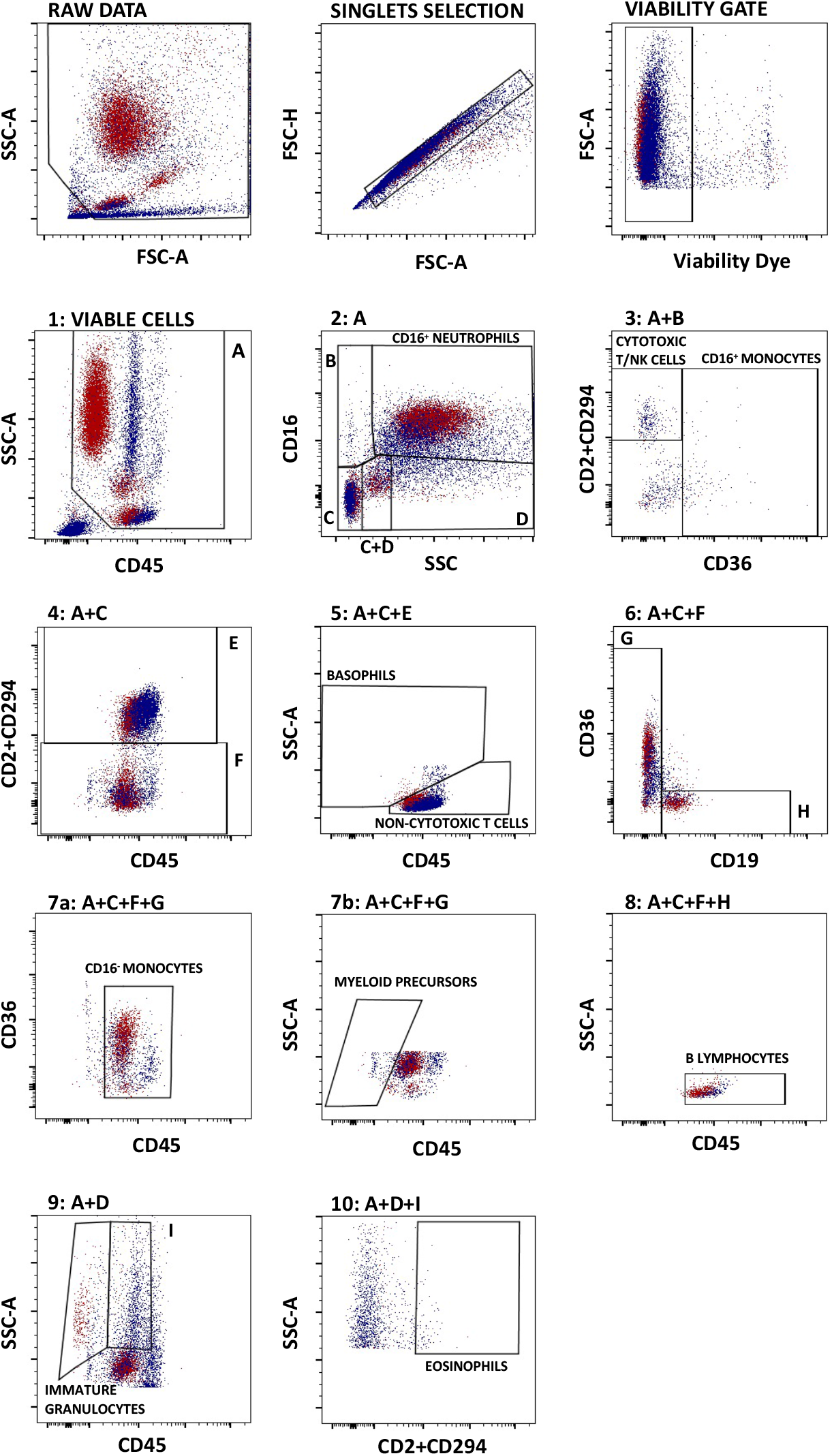
Typical overlay plots (red: peripheral blood, blue: colostrum) detailing the gating strategy applied to identify 10 leukocyte subpopulations from peripheral blood and colostrum with original gates set from a blood sample. The data presented were obtained from one representative donor from the lean cohort.

**Table 2.**
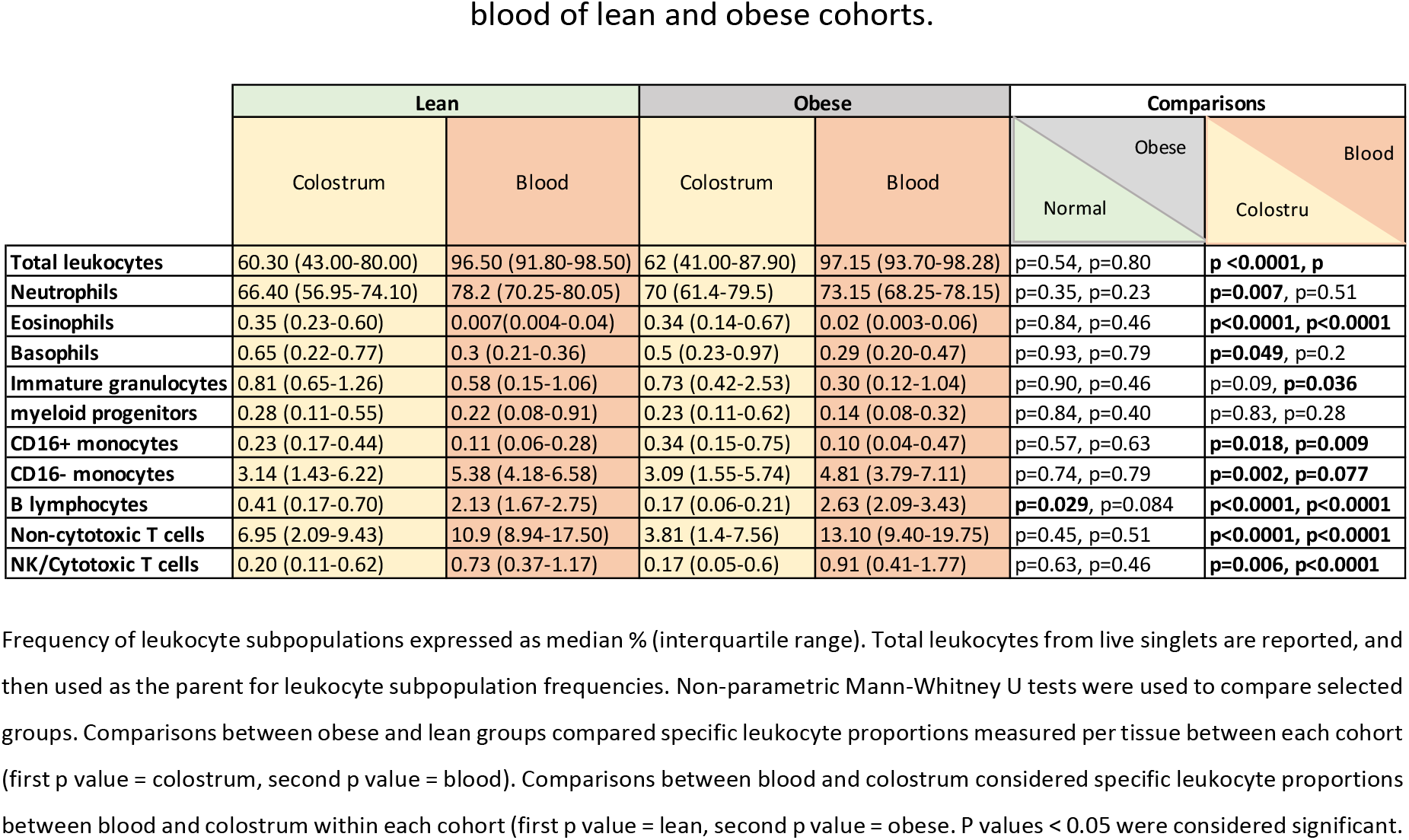
Summary of median leukocyte subset percentages identified in colostrum and peripheral blood of lean and obese cohorts.

Leukocytes comprised a significantly larger fraction of blood samples compared to colostrum samples, regardless of the study group (p<0.0001). While CD45^+^ leukocytes make up the large majority of nucleated blood circulating cells, rare CD45^-^ cells such as erythroid precursors or CD45^-^ megakaryocytes were recently reported in healthy individuals which may explain the < 3.5% CD45^-^ fraction identified in these samples^27,28^. Colostrum samples systematically exhibited a lesser leukocyte fraction. Colostrum has been found to contain 13.2-70.4% leukocytes, with epithelial cells, lactocytes, and progenitor and stem cells forming the non-leukocytic fraction^6,29^.

Overall, this panel classified > 78% and > 98% of CD45^+^ cells found in colostrum and peripheral blood of mothers, respectively. Therefore a small % of CD45^+^ cells present in colostrum remain to be identified, as shown by events remaining outside of population-calling gates (Figure 1, panels 6-10), consistent with the literature^7^. Neutrophils constituted the main leukocyte population present in both tissue types, as expected^7,30^. Neutrophil average proportions in colostrum were 1.5 to 5 times higher than previously reported in human colostrum using flow cytometry (medians: 66.40% and 70% in lean and obese groups, respectively, versus less than 15% in the study by Trend and colleagues), but similar to previously measured in colostrum using a blood hematology analyzer^7,31^. Blood circulating neutrophils have a lifespan of a few hours only, which is significantly shorter compared to other leukocytes^32^. Reducing the time between collection and analysis to < 3 h may have allowed increased detection of live neutrophils, compared to longer wait periods in other studies. Proportions measured in blood (medians: 78.20% and 73.15% in lean and obese groups, respectively) were higher than expected in peripheral blood, which is consistent with the literature reporting leukocytosis and impaired neutrophil apoptosis during pregnancy and labor^33^. There was no difference in neutrophil proportions in either tissue between both groups. However, there was significantly less neutrophils in lean mothers’ colostrum compared to lean mothers’ blood, while this difference was not recapitulated in the obese cohort (Table 2).

The second highest frequency of leukocytes in colostrum comprised non-cytotoxic T cells (medians: 6.95% and 3.81% in lean and obese groups, respectively, Table 2). While these proportions did not differ between cohorts, the frequency of non-cytotoxic T cells, described as CD45^+^SSC^Int^CD16^-^ CD2^+^CD294^+^ in peripheral blood was significantly higher compared to frequencies in colostrum, irrespective of the cohort (Fig. 2, Table 2). In contrast, previous work has described similar proportions of non-cytotoxic T cells in both tissues, although identified using a distinct gating strategy, as CD3^+^CD4^+^ events^31^.

**Fig 2.**
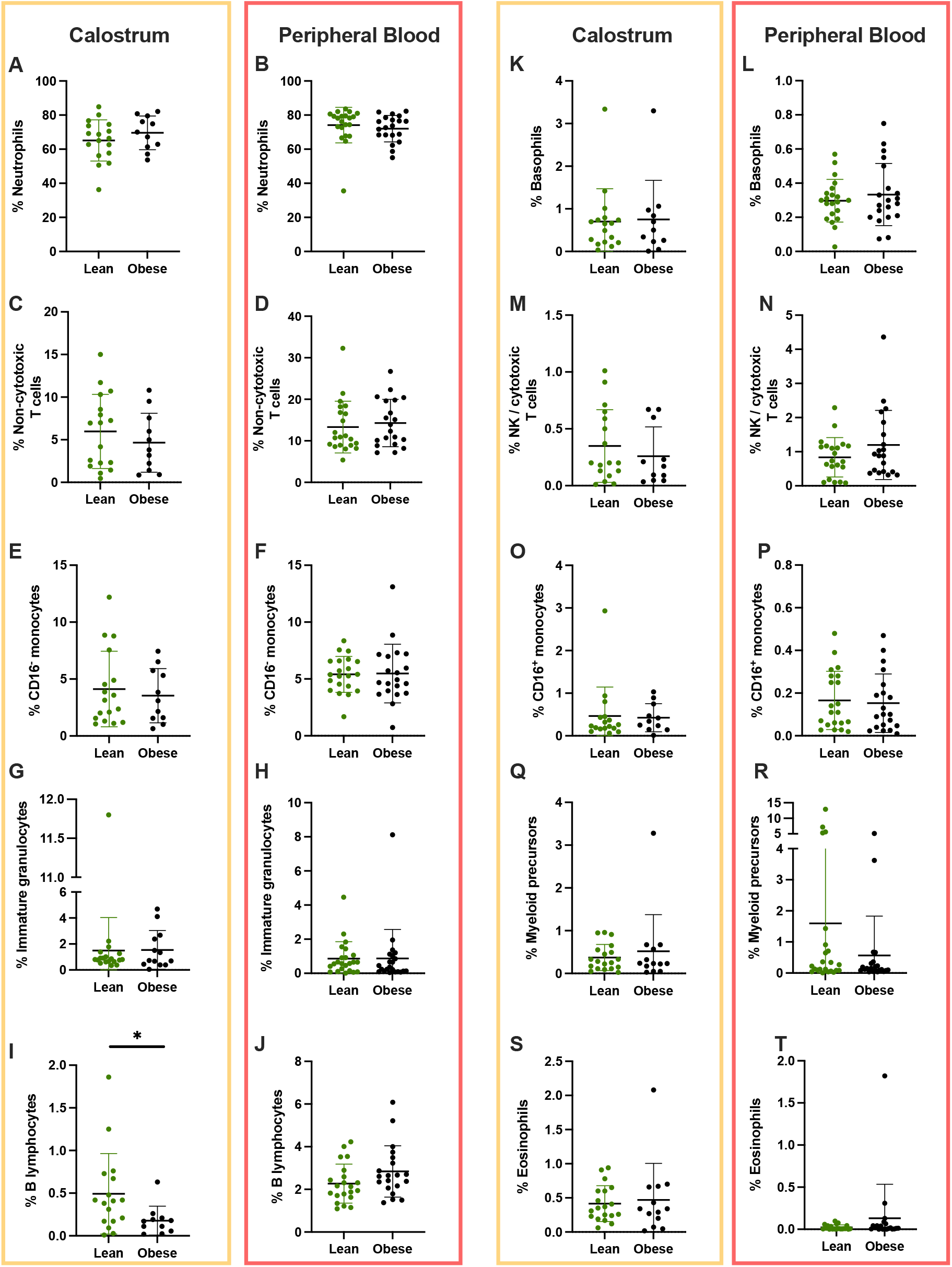
Relative frequencies of leukocyte subpopulations identified in colostrum and peripheral blood of lean and obese mothers.

Proportions of leukocytes subpopulations depended on tissue type. Overall, there was a significantly higher fraction of total leukocytes, CD16^-^ monocytes, B lymphocytes, non-cytotoxic T cells in peripheral blood compared to colostrum. On the other hand, we evidenced a significantly higher fraction of eosinophils and CD16^+^monocytes in colostrum compared to peripheral blood (Table 2). These trends were pervasive across study groups. Group-specific differences between tissue were also identified, with a significantly larger fraction of basophils and eosinophils in blood, solely in the lean group, and a significantly larger proportion of immature granulocytes in colostrum samples from the obese group only.

### 2.4 Major phenotypic changes are observed in leukocyte subpopulations between investigated tissues

Overlay of blood and colostrum samples from individual donors consistently highlighted various contrasts between the tissue types (Fig. 1). These included distinct internal complexity for the main 3 leukocyte populations as characterized by SSC, altered CD45 expression levels (Fig. 1, viable cells), and heterogenous CD16 expression on neutrophils (Fig. 1A). Therefore, we quantified these tissue-specific variations for each identified leukocyte population.

The relative size of cells within various leukocyte subpopulations varied significantly between tissues as measured by forward light scattering (FSC). B lymphocytes were significantly larger in size in colostrum compared to peripheral blood. Larger B lymphocytes have been described in breastmilk, which has been explained by their differentiation towards plasma cells in this tissue. These larger B cells are also primed to secrete antibodies^34^. The diameter of B cells is known to increase during infections as lymphoblastoid cells emerge^35^. Given the variety of microorganisms encountered in the breastmilk microbiota, it is not surprising that antigen-secreting cells be present in this tissue. On the other hand, basophils, both identified populations of T lymphocytes, and both subpopulations of monocytes were significantly smaller in colostrum compared to blood (Fig. 3A). Decrease in cell size is traditionally linked to apoptosis and cell death, as these processes provoke chromatin condensation, reorganization of the cytoskeleton and eventually fragmentation into apoptotic bodies^36^. Although the applied staining panel contained a viability marker and focused on analysis of live cells only, the present study did not include an apoptosis marker, which may have helped resolve the cell shrinkage observed in most cell types in colostrum. Further, the monocyte compartment was comprised of significantly larger cells in blood compared to colostrum, with classical CD16^-^ monocytes significantly larger in size compared to non-classical CD16^+^ monocytes, and this difference was recapitulated in colostrum (Fig. 3A). CD16^-^ monocytes are less susceptible to apoptosis compared to CD16^+^ monocytes, which could explain the size difference observed^37^. However, small and large subpopulations of monocytes have recently been described in peripheral blood in healthy state, with classical CD16^-^ monocytes comprising the majority of populations of both sizes, and cell-size dependent responses to stimulation within donors, which supports size differences being an actual phenotype unlinked to cell death^38^. In addition, the neutrophil fraction, which is the most delicate leukocyte cell subtype with high sensitivity to shear stress and temperature changes, did not present a significant variation in size between tissues, casting doubt on apoptosis being the cause for all observed cell shrinkage^39,40^. Differential isotonic pressure of tissue significantly impacts the size of leukocytes, which may be an alternative explanation for size differences observed as breastmilk has higher osmolality than peripheral blood^41,42^.

**Fig.3.**
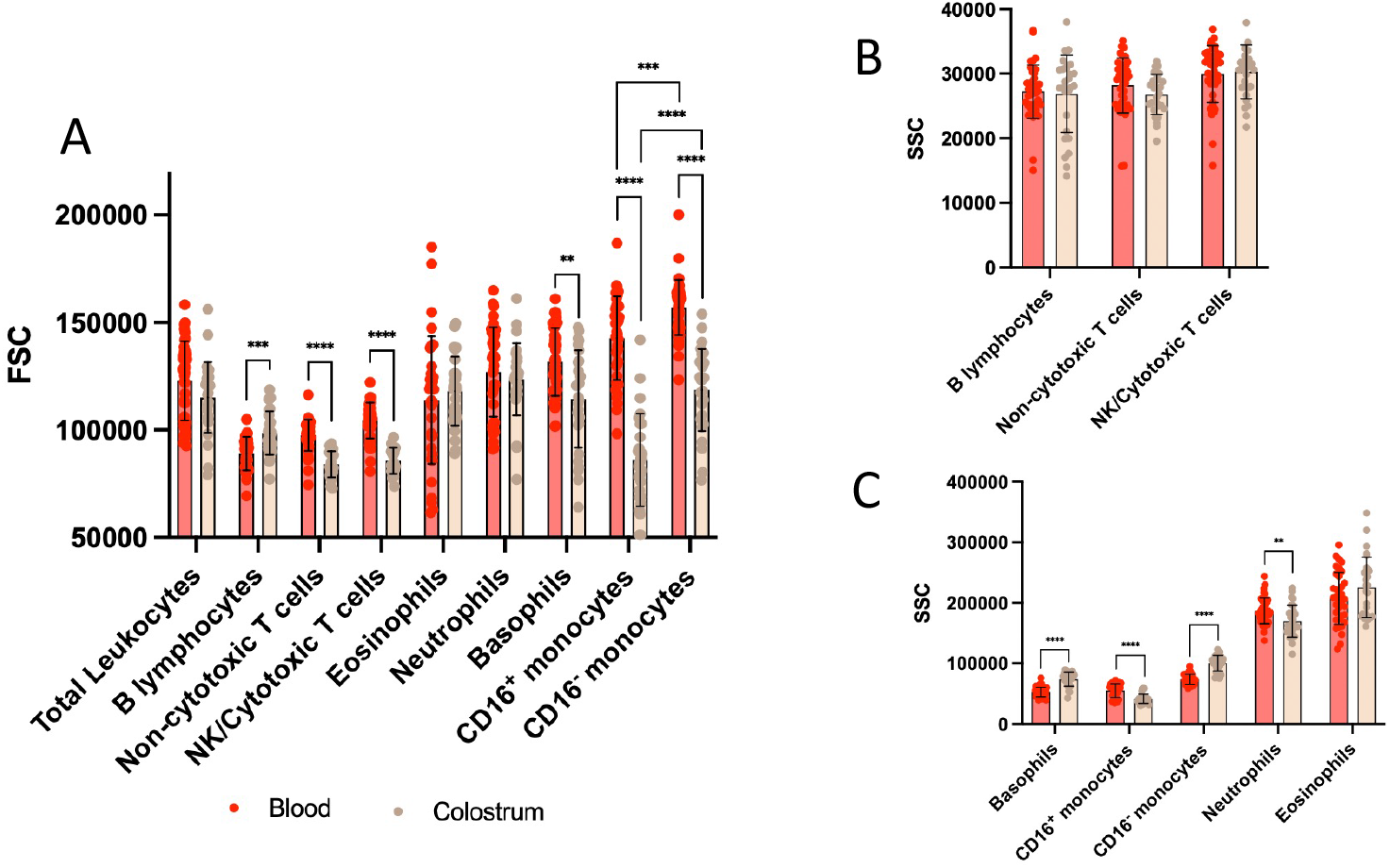
Relative size (FSC) and internal complexity (SSC) of identified leukocyte subpopulations in blood (red) and colostrum (beige) samples. p<0.05 (*), p<0.01 (**), p<0.001 (***), p<0.0001(****).

There was a cell-specific regulation of internal complexity, with lymphoid cells showing constant light side scattering (SSC), regardless of their origin, while SSC of myeloid cells were significantly regulated between tissues (Fig. 3B-C). Basophils and classical CD16^-^ monocytes were more internally complex in colostrum, while neutrophils and non-classical CD16^+^ monocytes were more internally complex in peripheral blood. As monocytes and neutrophils contain cytoplasmic granules, it is possible that some degranulation occurs in colostrum, known to contain a rich microbiota that may activate immune cells^43,44^. The increased internal complexity measured in colostrum for basophils and classical monocytes remains a matter of debate. Particularly, the contrast in SSC between both monocyte populations between tissues is intriguing, and recapitulates differences observed in colostrum FSC (Fig. 3A). Future transcriptional studies and detailed proteomic analyses may help elucidate the responsible mechanisms and resulting immunological phenotypes.

### 2.5 CD45 and CD16 expressions are regulated between tissues

After identifying distinct FSC and SSC profiles of leukocyte subpopulations depending on the tissue of origin, we asked whether relevant surface proteins were similarly regulated between peripheral blood and colostrum. CD45 is a transmembrane protein present on all mature leukocytes except for plasma cells and erythrocytes. As a protein tyrosine phosphatase, CD45 is involved in signal transduction in hematopoiesis, participates in antigen-mediated signaling in lymphocytes, and has been linked to neutrophil activation and migration^45^.

Overall, the relative CD45 expression was higher on lymphocytes and monocytes and lower on granulocytes, as expected^46^. There was no difference across tissues in the relative expression of CD45 on the surface of cells from the lymphoid lineage (Fig. 4A). Increased B lymphocyte size in colostrum suggested a differentiation towards plasma cells in this tissue. However we did not observe a decrease in CD45 expression on the surface of colostrum B cells, as expected for plasma cells^47^. Differentiating plasma cells progressively loose various surface markers including CD45 but also CD19, such that differentiated plasma cells are not included in the “B lymphocyte” compartment presented here, while no CD45 or CD19 loss is expected at the plasmablast stage, fitting with larger cell size findings of this study.

**Fig. 4.**
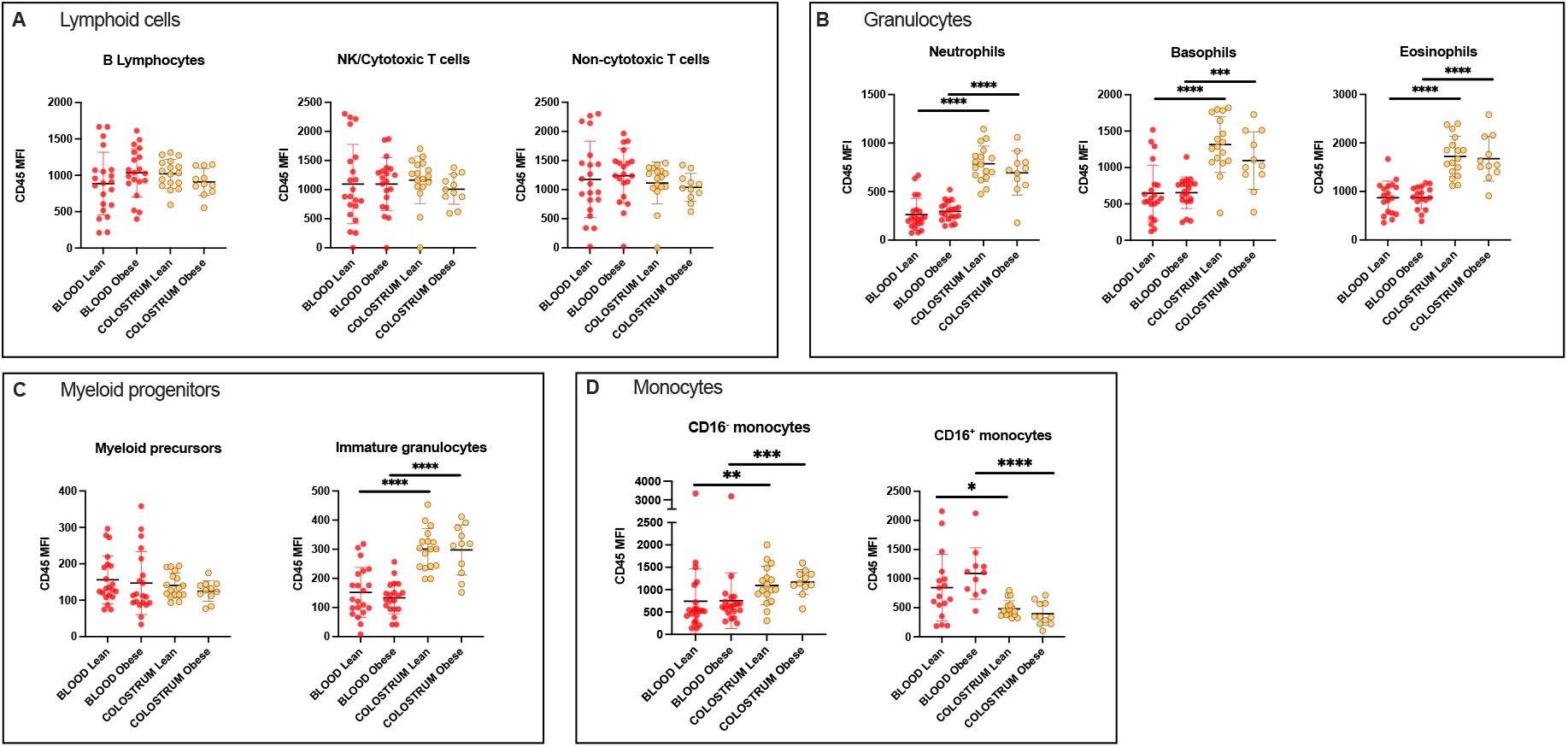
Relative expression of CD45 expressed as Median Fluorescence Intensity (MFI), on the surface of leukocytes from peripheral blood (red) or colostrum (beige) samples from obese and lean cohorts. Individual values are plotted together with mean and SD. p<0.05 (*), p<0.01 (**), p<0.001 (***), p<0.0001(****).

On the other hand, all 3 granulocyte subtypes exhibited a very significant increase in CD45 expression in colostrum compared to blood (Fig. 4B). CD45 upregulation has been described on granulocytes upon exposure to pathogenic microbes and physiological activators such as fMLP^48-50^. However, the implications of this regulation on the development of the immune response remain unclear, and conflicting results have been described. For example in neutrophils, upregulation of CD45 is consistent with their activation^50^. CD45 is also partially involved in regulating various neutrophil immune functions like cell adhesion, phagocytosis and ROS production^51^. However CD45 was also shown to downregulate neutrophil chemotaxis, and neutrophil ROS production in turn was shown to inhibit CD45^52,53^. Therefore, the present results warrant future in-depth analyses of the activation status of granulocytes present in colostrum using functional assays. Further, while early myeloid precursors did not exhibit changes in CD45 expression, with Median Fluorescence Intensity (MFI) consistently averaging around 150 in all groups, colostrum immature granulocytes had twice the levels of CD45 expression observed on the same population in peripheral blood, with MFI averaging from 134 and 128 in blood to >300 in colostrum, irrespective of the study groups (Fig. 4C). CD45 expression progressively increases during granulocytic maturation^54^. The sharp increase in CD45 expression on colostrum immature granulocytes may be the consequence of interactions with colostrum microbiota, as patients with acute infections of various origins exhibit increase of CD45RA on neutrophil surface which also correlate with increased frequency of circulating band, less mature neutrophils^55^.

Both monocyte subpopulations presented similar levels of CD45 in peripheral blood, with nonsignificant variations (CD16^+^ monocytes: obese cohort median MFI = 1107, vs. lean cohort median MFI = 697; CD16^-^ monocytes: obese cohort median MFI = 519, vs. lean cohort median MFI = 628, Fig. 4D). However, there was a significant regulation of CD45 expression on colostrum cells, and surprisingly the direction of this regulation was dictated by the subtype of monocyte. Classical CD16^-^ monocytes exhibited a significant increase in CD45 expression from blood to colostrum, while non-classical CD16^+^ monocytes experienced a significant downregulation of CD45 between blood and colostrum. Both trends were recapitulated across groups but were more significant in the obese mothers cohort (Fig. 4D).

CD16 (FcγRIII) is a transmembrane receptor for the crystallizable fraction of IgG antibodies and its crosslinking is involved in a variety of cell processes downstream of activation, such as ADCC in NK cells and degranulation and antibody-mediated phagocytosis in neutrophils. We observed consistent levels of CD16 on cytotoxic and NK cells, and non-classical CD16^+^ monocytes, across all study groups (Fig. 5A and C). On the other hand, there was a significant downregulation of CD16 on the surface of colostrum neutrophils compared to blood, irrespective of the study group. Colostrum B lymphocytes also exhibited a significant decrease in CD16 expression in the lean group only (mean MFI in lean group blood = 38.1 vs. mean MFI in lean group colostrum = 27.1, p=0.013). As CD16 is downregulated on plasma cells, it is possible that the smaller B cell fraction in obese group colostrum is accompanied by phenotypic disfunction evidenced from the absence of CD16 downregulation during differentiation towards plasma cells^56^. Downregulation of CD16 on neutrophils correlates with various events such as chemotaxis, activation and apoptosis^57,58^. Together with consistent size and increased CD45 expression in colostrum, these results do not support apoptosis of neutrophils in colostrum, but future experiments should investigate neutrophil apoptosis and activation status in this tissue.

**Fig. 5.**
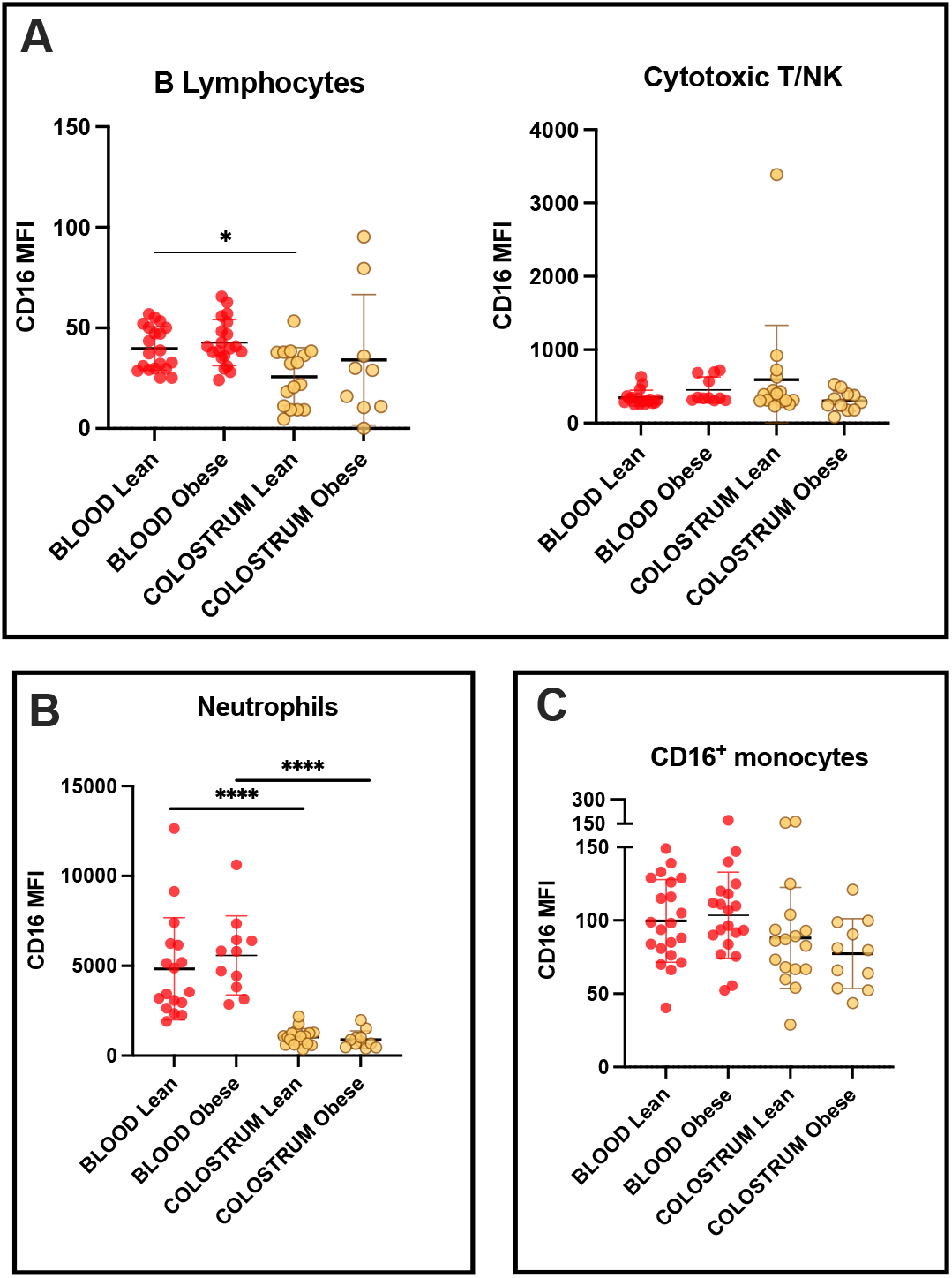
Relative expression of CD16 expressed as MFI on the surface of relevant leukocyte subpopulations contained in peripheral blood (red) or colostrum (beige) samples from obese and lean cohorts. Individual values are plotted together with mean and SD. p<0.05 (*), p<0.0001(****).

### 2.3 Obesity is associated with a smaller fraction of B lymphocytes in colostrum

A major finding of this study was evidencing a significantly reduced fraction of B lymphocytes in obese mothers’ colostrum compared to lean mothers’ colostrum (Fig.2 and Table 2, medians 0.17% vs. 0.41%, respectively, p = 0.029). On the other hand, in peripheral blood, B lymphocyte proportions were within similar ranges for both cohorts (medians: 2.13% and 2.63%), which is consistent with the literature using a similar gating strategy^59^. Comparing tissue types, the B cell compartment was systematically larger in blood compared to colostrum (Fig. 3A). This difference was statistically significant and consistent with earlier work^31^.

## 3. DISCUSSION

To the best of our knowledge, this is the first comprehensive study of the main leukocyte subtypes in colostrum from a Latin-American Mestizo population, the first report of phenotypic alterations of leukocyte subpopulations between peripheral blood and colostrum, and the first evidence of obesity altering colostrum leukocytes ^20^.

This work provides insights into the regulation of leukocyte trafficking from blood to colostrum since various significant trends were equally recapitulated in both mothers cohorts. Overall, the data indicate minimal regulation of the lymphoid compartment between tissues while myeloid cells were significantly altered morphologically and on the cell surface in colostrum. Notably, neutrophils size was constant between tissues, contrasting with the expected phenotype for apoptotic cells. Neutrophil CD45 levels were significantly upregulated in colostrum, while internal complexity was reduced, and CD16 downregulated. This may suggest activation, degranulation and CD16 shedding by tertiary granule-released metalloproteases, respectively, although further investigations are necessary^60^.

Obesity significantly reduced the B lymphocyte compartment in colostrum, without affecting peripheral blood. Unlike observed in the lean cohort, obese B lymphocytes did not exhibit a significant downregulation of CD16 from blood to colostrum. The relevance of this regulation remains to be investigated through a detailed characterization of B lymphocyte subpopulations, as evidence suggests peripheral blood B cells from obese individuals are more inflammatory and less efficient at switching to memory B cells upon antigen exposure^17^. Furthermore, as observed features of colostrum B cells hint toward antibody-secreting cells, these results suggest an impact of obesity on the quantity and quality of passive immunity provided to nursing infants.

As an exploratory study, this report highlights various key findings that require further study. First, what are the causes of the reduced B cell compartment in obese mothers’ colostrum, and what are the short- and long-term consequences in suckling infants? Why and how are leukocytes trafficked from blood to colostrum, and is the altered phenotype in colostrum a requisite for, or a consequence of trafficking? Finally, the presented data hint toward activation of the innate immune system in colostrum, accentuating the need to investigate colostrum as a complex system, together with its microbiota. Host-microbe crosstalk should be considered in future studies to shed light on the mechanistic regulation of colostrum composition in obesity, and its impact on suckling infants.

## 4. MATERIAL AND METHODS

### 4.1 Human samples

This study was approved by the Ethics Committee of the Hospital Regional Materno Infantil, Servicios de Salud de Nuevo León, Mexico, and the IRB at Escuela de Medicina y Ciencias de la Salud, TecSalud, in Monterrey, Mexico, with the ID CarMicrobioLHum-2018. All samples were collected and used following signed informed consent. Mothers provided blood and colostrum samples once, within 2 days of giving birth. Briefly, 3-4 ml of peripheral blood were collected in K_2_EDTA and placed on ice until processing. One-to-3 <u>mL</u>of colostrum per donor was obtained through pump-assisted milk extraction and immediately stored on ice, following washing of the breast and nipple area using soap and water. All samples were processed within 3 h of collection.

### 4.2 Leukocyte enrichment from colostrum

Around 1 ml of colostrum was processed for cell enrichment prior to staining for flow cytometry. Briefly, samples were centrifuged at 400 rcf for 15 min at 4°C. The supernatant was discarded, and the cell pellet was washed twice with PBS/2% FBS. Cells were counted and aliquoted for flow cytometry staining.

### 4.3 Colostrum-enriched cells staining

Depending on availability, between 200,000 and 1 x10^6^ cells were used for staining, and the same number of cells per sample were kept as unstained control. The same antibody lots were used to stain both types of tissues and antibody titration optimizations were performed for each tissue type to optimize resolution of fluorescence intensity over background. Cells were resuspended in the antibody master mix, which consisted of 2.5 μL mouse anti-human CD2-APC (BD^®^ cat. 560642), 5 μL mouse anti-human CD16-APC-H7 (BD^®^ cat. 560195), 5 μL mouse anti-human CD19-V450 (BD^®^ cat. 560353), 2.5 μL mouse anti-human CD36-PE (BD^®^ cat. 555455), 5 μL mouse anti-human CD45-V500 (BD^®^ cat. 560777), and 1.25 μL rat anti-human CD294-Alexa Fluor 647 (BD^®^ cat. 558042), in a final 100 μL staining volume with PBS + 2% FBS per 10^6^ cells. Samples were then incubated for 30 min on ice in the dark, then washed and resuspended in PBS/2% FBS. Ten min before acquisition, propidium iodide (BD^®^ cat. 556463) was added to the tube as per manufacturer’s recommendations.

### 4.4 Peripheral blood staining

Fifty μL anticoagulated peripheral blood were stained using 2.5 μL CD2-APC, 1.25 μL CD16-APC-H7, 5μL CD19-V450, 2.5 μL CD36-PE, 1.25 μL CD45-V500, and 1.25 μL CD294-Alexa Fluor 647, in a final 100 μL staining volume with PBS + 2% FBS for 30 minutes on ice, in the dark. Samples were then subjected to erythrocyte lysis using BD^®^ Pharm Lyse solution (BD^®^ cat. 555899) as per manufacturer’s instructions, or 500 μL of 1.5 M NH_4_Cl solution for 10 min at room temperature, followed by immediate washing and resuspension in PBS/2% FBS. Ten min before acquisition propidium iodide was added to tubes as per manufacturer’s recommendations.

### 4.5 Flow cytometry data acquisition

Samples were analyzed on a BD^®^ FACSCelesta flow cytometer fitted with 405 nm, 488 nm, and 633 nm lasers and operated through the BD^®^ FACSDiva software v.8. Compensation controls were prepared using compensation beads (Anti-mouse Ig,K Neg Control compensation, BD^®^ cat. 552843) following manufacturer’s recommendations. At least 30,000 events were recorded from every sample, with FSC event threshold adjusted to 35,000 for peripheral blood and 28,000 for colostrum samples.

### 4.6 Analysis and gating strategy

Cytometry data were analyzed using FlowJo software v.10 (Treestar LLC). Automatic compensation was performed prior to analysis. A strict quality control workflow was established, where samples had to exhibit a characteristic FSC/SSC morphology, viability above 85% from singlet gate, and >10,000 leukocytes (CD45^+^ cells) acquired, to be included in the final analysis and comparisons. The gating strategy applied has been previously described^7,21^.

### 4.7 Statistical Analysis

Proportions of leukocyte subsets were analyzed using mean and standard deviation. Means of Median Fluorescence Intensity (MFI) of a surface marker’s expression were compared using Mann-Whitney tests using GraphPad Prism v. 9. Graphs are showing discrete data and mean with SD.

## 5. AUTHORSHIP

R.P.S processed the samples, analyzed the data, and wrote the manuscript. J.A.C.C processed the samples, analyzed the data, and reviewed the manuscript. V.J.L.D., J.M.V and M. R. A. G. organized and conducted the human study, acquired samples, and edited the manuscript. E.V.G. analyzed the data and wrote the manuscript. C.L.C designed the study, analyzed the data, and edited the manuscript. V.L. and G.G.R. participated in study design, data analysis, and reviewed the manuscript. M.B. designed the study, analyzed the data, wrote, and edited the manuscript.

## 6. ACKNOWLEDGMENTS

This research was supported by Tecnológico de Monterrey and TecSalud, and student scholarships from the National Research Council CONACYT of Mexico (743743 and 1007842) We also thank StrainBiotech SAPI de C.V. for partial financing this work.

